# Mating-type specific ribosomal proteins control aspects of sexual reproduction in *Cryptococcus neoformans*

**DOI:** 10.1101/777102

**Authors:** Giuseppe Ianiri, Yufeng “Francis” Fang, Tim A. Dahlmann, Shelly Applen Clancey, Guilhem Janbon, Ulrich Kück, Joseph Heitman

## Abstract

The *MAT* locus of *Cryptococcus neoformans* has a bipolar organization characterized by an unusually large structure, spanning over 100 kb. *MAT* genes have been characterized by functional genetics as being involved in sexual reproduction and virulence. However, classical gene replacement failed to achieve mutants for five *MAT* genes (*RPL22, RPO41, MYO2, PRT1, RPL39*), indicating that they are likely essential. In the present study, targeted gene replacement was performed in a diploid strain for both the α and **a** alleles of the ribosomal genes *RPL22* and *RPL39*. Mendelian analysis of the progeny confirmed that both *RPL22* and *RPL39* are essential for viability. Ectopic integration of the *RPL22* allele of opposite *MAT* identity in the heterozygous *RPL22***a/***rpl22*αΔ or *RPL22*α*/rpl22***a**Δ mutant strains failed to complement their essential phenotype. Evidence suggests that this is due to differential expression of the *RPL22* genes, and an RNAi-dependent mechanism that contributes to control *RPL22***a** expression. Furthermore, via CRISPR/Cas9 technology the *RPL22* alleles were exchanged in haploid *MAT*α and *MAT***a** strains of *C. neoformans*. These *RPL22* exchange strains displayed morphological and genetic defects during bilateral mating. These results contribute to elucidate functions of *C. neoformans* essential mating type genes that may constitute a type of imprinting system to promote inheritance of nuclei of both mating types.

## Introduction

Infectious diseases cause significant morbidity and mortality worldwide in both developed and developing countries. Fungal infections are common in humans and impact the majority of the world’s population, but are often underestimated (BROWN *et al.* 2012). The *Cryptococcus* species complex includes basidiomycetous fungal pathogens that can cause lung infections and life-threatening meningoencephalitis in both normal and immunocompromised patients, accounting for approximately 1 million annual infections globally and almost 200,000 annual mortalities (RAJASINGHAM *et al.* 2017). The available drugs to treat *Cryptococcus* infections are amphotericin B, 5-flucytosine, and azoles. These drugs are characterized by limited spectrum, toxicity, unavailability in some countries, and emergence of drug resistance (BROWN *et al.* 2012).

In fungi, the mechanisms that govern sexual reproduction are controlled by specialized regions called mating type (*MAT*) loci. The genomic organization of the *MAT* loci can differ among fungi. The tetrapolar mating type includes two *MAT* loci, the P/R locus encoding pheromones and pheromone receptor genes defining sexual identity and mate recognition, and the HD locus encoding homeodomain transcription factors that govern post-mating developmental processes. In the tetrapolar system the P/R and HD loci are located on different chromosomes and segregate independently during meiosis, hence generating recombinant *MAT* systems. Conversely, in the bipolar mating system both the P/R and the HD loci are linked on the same chromosome, and recombination in this region is suppressed. A variant of the bipolar system is a mating conformation called pseudobipolar, in which the P/R and the HD loci are located on the same chromosome but unlinked, thus allowing (limited) recombination (COELHO *et al.* 2017).

*C. neoformans* has a well-defined sexual cycle that is controlled by a bipolar *MAT* system that is derived from an ancestral tetrapolar state. The *C. neoformans MAT* locus evolved in a unique configuration as it spans over 100 kb and contains more than 20 genes that control cell identity, sexual reproduction, infectious spore production, and virulence. The two opposite *C. neoformans MAT*α and *MAT***a** alleles include divergent sets of the same genes that evolved by extensive remodeling from common ancestral DNA regions. Both the *MAT*α and *MAT***a** allele contain five predicted essential genes: *RPO41, PRT1, MYO2, RPL39,* and *RPL22* (LENGELER *et al.* 2002; FRASER *et al.* 2004). Rpo41 is a mitochondrial RNA polymerase that transcribes mitochondrial genes and also synthesizes RNA primers for mitochondrial DNA replication (SANCHEZ-SANDOVAL *et al.* 2015). Prt1 is a subunit of the eukaryotic translation initiation factor 3 (eIF3) that plays a critical role in translation (BEZNOSKOVA *et al.* 2015). Myo2 is a myosin heavy chain type V that is involved in actin-based transport of cargos and is essential in *S. cerevisiae* (JOHNSTON *et al.* 1991). Rpl39 and Rpl22 are ribosomal proteins.

This study focused on the *MAT* ribosomal proteins, demonstrating that both *RPL39* and *RPL22* α and **a** alleles are essential in *C. neoformans.* Because Rpl22 in yeast and vertebrates has been found to play specialized functions and extra-ribosomal roles (GABUNILAS AND CHANFREAU 2016; KIM AND STRICH 2016; ZHANG *et al.* 2017; ABRHAMOVA *et al.* 2018), we aimed to characterize the functions of the *C. neoformans RPL22*α and *RPL22***a** genes. We found that ectopic integration of an *RPL22* allele failed to complement the essential phenotype due to the mutation of the *RPL22* allele of opposite mating type. We found differential expression of the *C. neoformans RPL22*α and *RPL22***a** genes during mating, and discovered an RNAi-mediated mechanism that contributes to control *RPL22***a** expression. Next, using the CRISPR/Cas9 technology, *RPL22* alleles were exchanged in haploid *MAT*α and *MAT***a** strains of *C. neoformans* and this resulted in morphological and genetic defects during bilateral mating. In summary, these studies reveal a novel role for diverged essential ribosomal proteins in controlling fungal sexual reproduction.

## Materials and methods

### Strains and culture conditions

The strains utilized in the present study are listed in table S1. Heterozygous mutants were generated in the diploid *C. neoformans* strain AI187 (*MAT*α/*MAT***a***ade2*/*ADE2 ura5*/*URA5*) according to a previously developed strategy (IANIRI AND IDNURM 2015). *C. neoformans* strain AI187 was generated through the fusion of strains JF99 (*MAT***a***ura5*) and M001 (*MAT*α *ade2*) (IDNURM 2010). For transformation of haploid *C. neoformans* strains, we employed H99α and KN99**a** (NIELSEN *et al.* 2003). All of the strains were maintained on yeast extract-peptone dextrose (YPD) agar medium.

### Molecular manipulation of *C. neoformans*

For the generation of heterozygous mutants, 1.5 kb regions flanking the genes of interest were amplified by PCR and fused with the *NAT* marker through *in vivo* recombination in *S. cerevisiae* as previously described (IANIRI AND IDNURM 2015). Split-marker gene replacement alleles were amplified from *S. cerevisiae* transformants with primers JOHE43263/ALID1229 and JOHE43264/ALID1230 in combination with ai37 and JOHE44324, respectively. The amplicons were precipitated onto gold beads and transformed into C. neoformans with a Bio-Rad particle delivery system (TOFFALETTI *et al.* 1993); W7 hydrochloride was added to YPD + 1 M Sorbitol to increase the efficiency of homologous recombination (ARRAS AND FRASER 2016). Transformants were selected on YPD + NAT and screened for homologous recombination events by PCR with primers external to the replaced regions in combination with primers specific for the *NAT* marker, and with gene-specific internal primers. The primers used are listed in Table S2.

For complementation experiments, a region of 2397 bp including the *RPL22*α gene with its promoter and terminator was amplified by PCR from *C. neoformans* H99α genome and cloned in pCR™2.1 according to the manufacturers’ instructions. Similarly, a region of 2648 bp including the *RPL22***a** gene with its promoter and terminator was amplified by PCR from *C. neoformans* KN99**a** genome and cloned in pCR™2.1 according to manufacturers’ instruction. Plasmids were recovered from *E. coli* TOP10 and sequenced to identify error-free clones (Table S2). Sequence confirmed plasmids were digested with SpeI-XhoI and SpeI-NotI to obtain regions including the *RPL22*α and *RPL22***a** genes, respectively. These fragments were purified and subcloned within the pSDMA57 plasmid for safe haven complementation (ARRAS *et al.* 2015) digested with the same enzymes. The recombinant plasmids were recovered from *E. coli* TOP10, linearized with AscI, PacI, or BaeI, and transformed through biolistic in *C. neoformans* heterozygous mutants GI56 (*RPL22*α*/rpl22***a**Δ) and GI81 (*RPL22***a/***rpl22*αΔ) as described above. *C. neoformans* transformants were selected on YPD + neomycin G418, and subjected to DNA extraction and PCR analyses to identify transformants having the correct insert within the safe haven region (ARRAS *et al.* 2015).

The recently-developed TRACE technology (Transient CRISPR/Cas9 coupled with electroporation) (FAN AND LIN 2018) was utilized for the generation of the 5ʹΔ *RPL22***a** strain GI228 and the *RPL22* exchange alleles. For 5ʹΔ *RPL22***a**, a homology directed repair (HDR) template consisting of 1.5 kb sections flanking the region upstream *RPL22***a** targeted by sRNA was fused with the *NAT* marker through *in vivo* recombination in *S. cerevisiae* as described above.

For the generation of the *RPL22* exchange alleles, we developed a dual CRISPR/Cas9 system to exchange the two different *RPL22* alleles alone, and insert selectable markers (*NAT* or *NEO*) separately in the Safe Haven 2 (*SH*2) region. HDR templates were generating by fusing ~1.0 kb fragments flanking the *RPL22* genes with the ORF of the opposite *RPL22* gene. For the generation of a chimeric c*RPL22*α (c = chimeric), the N-terminal region of *RPL22*α (from nucleotide 1 to 268) and the C terminal region of *RPL22***a** (from nucleotide 253 to 600) were combined together by PCR, fused with ~1.0 kb regions flanking the *RPL22***a** gene, and employed as an HDR template. All HDR templates were assembled using overlap PCR as described in (DAVIDSON *et al.* 2002).

Specific guide RNAs (gRNA) were designed according to (FANG *et al.* 2017) using EuPaGDT (http://grna.ctegd.uga.edu/) available on FungiDB (https://fungidb.org/fungidb/). Complete gRNAs were generated by one-step overlap PCR, in which a bridge primer that comprises the 20 nucleotide gRNA guide sequences was utilized to integrate the U6 promoters (amplified from *C. deneoformans* XL280 genomic DNA) and the gRNA scaffold [amplified from the plasmid pYF515 (FANG *et al.* 2017)]. *CAS9* was amplified from pXL1-Cas9 (FAN AND LIN 2018). Safe Haven 2 (SH2) sequence was obtained from (UPADHYA *et al.* 2017). All PCR-amplifications were conducted using Phusion High-Fidelity DNA Polymerase (NEB). *C. neoformans* was transformed with *CAS9*, gRNAs, and HDR templates in through electroporation following the previously reported protocol (FAN AND LIN 2018). Transformants were screened for homologous recombination events by PCR as previously indicated.

### Genetic analyses and scanning electron microscopy of reproductive structures

The heterozygous strains generated were grown on Murashige-Skoog (MS) medium to induce meiosis and sporulation. Haploid *C. neoformans* mutants were crossed with *C. neoformans* WT strains of compatible mating type (H99 *MAT*α and KN99**a***MAT***a**) on MS medium and monitored for the formation of sexual structures. Spores were micromanipulated and allowed to germinate onto YPD agar for 3 to 4 days at 30°C, and then tested for the segregation of the genetic markers. For the heterozygous strains the markers were nourseothricin resistance (NAT^R^) or sensitivity (NAT^S^), *ura5/URA5*, *ade2/ADE2, MAT*α/*MAT***a**, plus neomycin G418 resistance (NEO^R^) or sensitivity (NEO^S^) for the complementing strains. For the haploid strains they were either NAT^R^-NAT^S^ or NEO^R^-NEO^S^, and *MAT*α or *MAT***a**. The analyses were performed by spotting 2 μl of cell suspensions onto YPD + nourseothricin (100 μg/ml) or neomycin (100 μg/ml), YNB + adenine (20 mg/L) or YNB + uracil (40 mg/L). The mating type was scored by crossing haploid progeny to strains KN99**a** and H99 on MS media supplemented with adenine and uracil, and by evaluating the formation of sexual structures by microscopy (IDNURM 2010; IANIRI AND IDNURM 2015). For NAT^R^ colonies the mating type was confirmed by PCR with primers JOHE39201-JOHE39202 (*MAT***a**) and JOHE39203-JOHE39204 (*MAT*α).

For strain YFF116 (*rpl22***a**∷*RPL22*α *NEO*), genetic segregation of the *MAT* and *NEO* markers was carried out by crossing YFF116 x H99α on MS, and by dissecting recombinant progeny as described above. Progeny that germinated were subjected to 10-fold serial dilution on YPD, YPD + neomycin, and hydroxyurea (125 mM). To evaluate the consequences of the *rpl22***a**∷*RPL22*α genetic modification in unilateral and bilateral mating without the influence of the *NEO* marker, NEO^S^*MAT***a** progeny were crossed both with H99α and the YFF92 strain.

Scanning electron microscopy (SEM) was performed at the North Carolina State University Center for Electron Microscopy, Raleigh, NC, USA. Samples were prepared for SEM as previously described (FU AND HEITMAN 2017). Briefly, a small MS agar block containing hyphae was excised and fixed in 0.1 M sodium cacodylate buffer, pH = 6.8, containing 3% glutaraldehyde at 4°C for several weeks. Before imaging, the agar block was rinsed with cold 0.1 M sodium cacodylate buffer, pH = 6.8 three times and post-fixed in 2% osmium tetroxide in cold 0.1 M cacodylate buffer, pH = 6.8 for 2.5 hours at 4°C. Then the block was critical-point dried with liquid CO_2_ and sputter coated with 50 Å of gold/palladium with a Hummer 6.2 sputter coater (Anatech). The samples were viewed at 15KV with a JSM 5900LV scanning electron microscope (JEOL) and captured with a Digital Scan Generator (JEOL) image acquisition system.

### RT-qPCR analysis during mating and statistical analysis

For RT-qPCR analysis of *RPL22* expression during mating, strains were grown overnight in liquid YPD, and cellular density was adjusted to 1 × 10^9^ CFU/mL. Equal amounts of each cellular suspension of strains to be analyzed were mixed, and 5 spots of 300 μl were placed onto one plate of MS agar per day of incubation. Control conditions were the single strains on YPD agar (2 spots of 300 μl per day of incubation). Every 24 h, cells were scraped off the MS plate, washed once with sterile water, lyophilized, and kept at −80°C until RNA extraction. RNA extraction was performed with the standard TRIzol protocol following the manufacturers’ instructions (RIO *et al.* 2010). Extracted RNA was treated with DNase and purified with an RNA clean and concentration kit (Zymo Research). Then, 3 μg of purified RNA were converted into cDNA via the Affinity Script QPCR cDNA synthesis kit (Agilent Technologies). cDNA synthesized without the RT/RNase block enzyme mixture was utilized as a control for genomic DNA contamination. Approximately 500 pg of cDNA were utilized to measure the relative expression level of target genes through quantitative real-time PCR (RT-qPCR) using the Brilliant III ultra-fast SYBR green QPCR mix (Agilent Technologies) in an Applied Biosystems 7500 Real-Time PCR System. A control without template RNA was included for each target. Technical triplicates and biological triplicates were performed for each sample. Gene expression levels were normalized using the endogenous reference gene *GDP1* and determined using the comparative ΔΔCt method.

To determine whether the relative gene expression levels between strains of the same mating reaction in the same day of incubation (for example, *RPL22*α and *RPL22***a** expression in WT H99α x KN99**a** cross after 48 h of incubation) exhibited statistically significant differences (p<0.05, p<0.01, p<0.001), the unpaired student’s t-test with Welch’s correction was applied. To compare the results of different strains in different mating reactions, ordinary one-way ANOVA with Tukey’s multiple comparison test was applied. Because in these comparisons we were interested in monitoring the changes in gene expression following genetic manipulation, only statistically significant differences (p<0.05, p<0.01, p<0.001) were displayed for the expression levels of the same gene on the same day of incubation in separate mating reactions (for example, *RPL22***a** expression in WT H99α x KN99**a** cross compared to *RPL22***a** expression in *rdp1*Δ x *rdp1*Δ bilateral cross after 48 h of incubation). Statistical analyses were performed using the software PRISM8 (GraphPad, https://www.graphpad.com/scientific-software/prism/).

### RNA structure modeling

RNA structure modeling was conducted with RNAfold (LORENZ *et al.* 2011) with default settings.

### sRNA data processing

Small RNA (sRNA) sequencing libraries from *C. neoformans* WT H99 × KN99**a** cross and *rdp1*Δ bilateral cross are as described in (WANG *et al.* 2010). The adapters sequences (5-prime: GTTCAGAGTTCTACAGTCCGACGATC; 3-prime: TCGTATGCCGTCTTCTGCTTGT) were removed by using cutadapt v1.9 (MARTIN 2011) and trimmed reads were mapped with bowtie v1.2.2 (LANGMEAD *et al.* 2009) against the *MAT***a** (AF542528.2) and *MAT*α (AF542529.2) loci from *C. neoformans* strains 125.91 and H99, respectively (LENGELER *et al.* 2000). Mapping was performed by allowing a single nucleotide mismatch and up to five alignments within both mating type loci, and the H99α genome. Furthermore, reads showing a single perfect match were considered in order to identify their genetic origin. Read counts were calculated with SAMtools’ depth function and by using custom made Perl scripts (LI *et al.* 2009; DAHLMANN AND KÜCK 2015). The read counts were normalized against tRNA mapping reads (tRNA read counts per 100,000 reads). The normalization factors for the WT mating and the *rdp1*Δ mating were calculated with 1.386 and 0.728, respectively.

### Chemical genetic screen and phenotypic analysis

Phenotypic analysis was performed for all the strains listed in table S1 with the standard 10-fold serial dilution method. Tested conditions and stresses included: temperatures of 4°C, 25°C, 30°C, 37°C, 38°C, 39°C; antifungal drugs, such as amphotericin B (AmB, 1.5 μg/mL), 5-fluorocytosine (5-FC, 100 μg/mL), fluconazole (FLC, 20 μg/mL), FK506 (1 μg/mL), rapamycin (1 μg/mL); cell wall and plasma membrane stressors, such as YPD and YP supplemented with NaCl (1.5 M and 1 M, respectively) and sorbitol (2M and 1.5 M, respectively), caffeine (10 mM), calcofluor white (4 mg/mL), Congo Red (0.8%); genotoxic, oxidative, nitrosative and other stress-inducing agents, such as ethidium bromide (10 μg/mL), sodium nitrite (NaNO_2_, 1.5 mM), UV (150 μJ x cm^2^), hydrogen peroxide (H_2_O_2_, 3 mM), cycloeximide (0.15 μg/mL), dithiothreitol (DTT, 15 mM), hydroxyurea (125 mM), tunicamycin (0.7 μg/mL), benomyl (2.5 μg/mL), cadmium sulphate (CdSO_4_, 30 μM). Unless indicated, plates were incubated at 30°C for 3 to 6 days and photographed.

### Data Availability Statement

Strains and plasmids are available upon request. The authors affirm that all data necessary for confirming the conclusions of the article are present within the article, figures, and tables.

## Results

### The *MAT* ribosomal genes *RPL22* and *RPL39* are essential

Both the α and **a** alleles of the five predicted *MAT* essential genes (*RPL39, RPL22, MYO2, RPO41, PRT1*) were identified in the genomes of *C. neoformans* H99α and KN99**a**, and were subjected to targeted mutagenesis in the *C. neoformans* diploid strain AI187 according to the strategy reported by Ianiri and Idnurm with minor modifications (IANIRI AND IDNURM 2015). Briefly, cassettes for targeted gene replacement were generated by *in vivo* recombination in *S. cerevisiae*, and amplified by PCR to perform targeted mutagenesis via split-marker coupled with the use of the non-homologous end joining (NHEJ) inhibitor W7 hydrochloride (FU *et al.* 2006; ARRAS AND FRASER 2016). These modifications were critical to increase the rate of homologous recombination, and allowed the generation of heterozygous mutants for both the *MAT*α and *MAT***a** alleles of the *RPL39*, *RPL22*, and *MYO2* genes. Despite these modifications, heterozygous mutants for *RPO41* and *PRT1* were not obtained. This study reports the genetic analysis of mutants for the *MAT* ribosomal genes *RPL39* and *RPL22*, and further focuses on the characterization of the *RPL22* gene.

Heterozygous mutants were confirmed by PCR analyses, and then transferred onto MS medium supplemented with adenine and uracil and allowed to undergo meiosis, sporulation, and basidiospore production. Spores were micromanipulated on YPD agar and subjected to phenotypic analysis to assess the segregation of the four available markers (*URA5, ADE2, MAT, NAT)*. Because the predicted essential genes were deleted by insertion of the *NAT* marker, the absence of NAT^R^ progeny indicates an essential gene function. The other 3 markers were tested to exclude defects in meiosis: while the *URA5/ura5* and *ADE2/ade2* loci were expected to segregate independently, the segregation of the *MAT* region was expected to be linked to the mutated alleles, with progeny being only *MAT***a** when derived from *MAT*α heterozygous deletion mutants, and only *MAT*α when derived from *MAT***a** heterozygous deletion mutants.

Heterozygous mutants for the *MAT* ribosomal proteins Rpl39 and Rpl22 produced basidiospores that displayed a rate of germination ranging from 36% to 41% (Table 1). Mendelian analysis of the progeny confirmed that both the *MAT***a** and *MAT*α alleles of Rpl39 and Rpl22 encode an essential function (Table 1). Figure 1 shows an example of the genetic analysis performed on progeny derived from strains GI233 (*RPL39***a**/*rpl39*αΔ) and GI56 (*RPL22*α*/rpl22***a**Δ). As expected, in all cases the progeny inherited only one *MAT* allele, which is the opposite of the mutated gene. One exception is that one NAT^R^ progeny was obtained from sporulation of the heterozygous *RPL22*α*/rpl22***a**Δ; further PCR analysis revealed that this strain has the mutated *rpl22***a**Δ*NAT* and an extra copy of the *RPL22*α gene, suggesting that its *NAT* resistance is likely due to aneuploidy of chromosome 5 (1n + 1) where the *MAT* locus resides.

**Table 1.** Genetic analysis of *C. neoformans* heterozygous mutants for the *MAT*α and *MAT***a** ribosomal proteins Rpl39 and Rpl22.

| Genotype | Strain | Protein function | Germinated basidiospores/<br>dissected basidiospores | NAT-R progeny/<br>basidiospore germinated | Phenotype |
| --- | --- | --- | --- | --- | --- |
| *RPL39α/rpl39aΔ* | GI230 | Large Subunit Ribosomal Protein L39 | 31/80 – 38.7% | 0/31 | Inviabile |
| *RPL39a/rpl39α $\Delta$* | GI233 |  | 29/70 – 41.4% | 0/29 | Inviabile |
| *RPL22α/rpl22aΔ* | GI56 | Large Subunit Ribosomal Protein L22 | 23/63 – 36.5% | 1/23* | Inviabile |
| *RPL22a/rpl22α $\Delta$* | GI81 |  | 28/72 – 38.8% | 0/28 | Inviabile |

\* The NAT-R progeny obtained is likely aneuploid for chromosome 5, because PCR confirmed the presence of the *RPL22 $\alpha$* allele and the mutated copy of the *RPL22a*.

**Figure 1.**
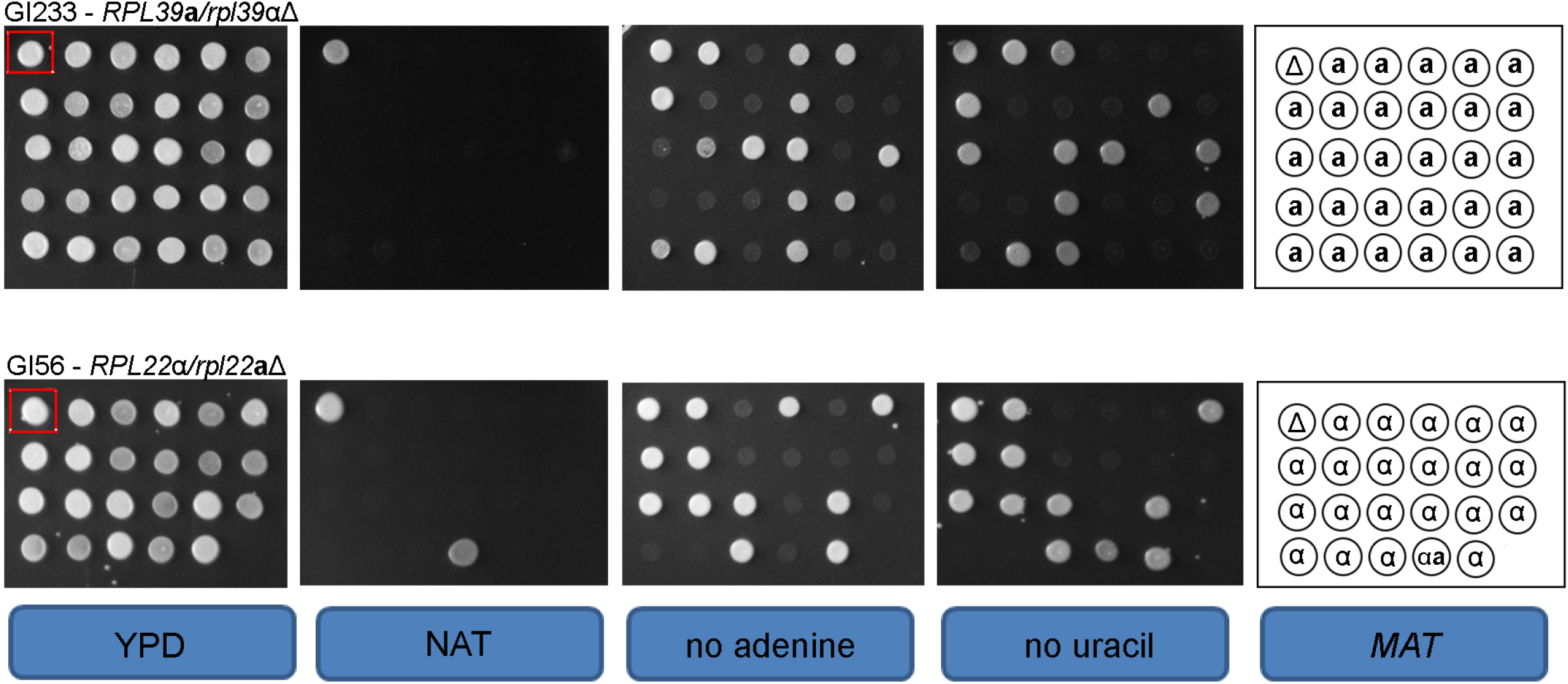
The *MAT* locus contains essential genes encoding ribosomal proteins Rpl39 and Rpl22. Representative example of genetic analysis of two *C. neoformans* heterozygous mutants [GI233 (*RPL39***a**/*rpl39*αΔ) and GI56 (*RPL22*α/*rpl22***a**Δ)]. The first colony in red box in the top left corner represents the original heterozygous mutant from which the progeny analyzed originated. The remaining isolates were haploid germinated spore progeny grown on control medium YPD, YPD + NAT, SD-uracil, and SD – adenine. The *MAT* type of the progeny is indicated on the right panel (Δ indicates the heterozygous mutant).

### The *C. neoformans RPL22* alleles are highly similar

The *RPL22***a** and *RPL22*α genes share 83% identity at the DNA level, and the encoded proteins differ in 5 amino acids that are located in the N-terminal region (Fig. S1 A, B). With the exception of intron 1 that is 132 bp for *RPL22*α and 116 bp for *RPL22***a**, both of the *C. neoformans RPL22* genes contain 4 exons and 3 introns of identical length. Intron 1 shares ~75% nucleotide identity, intron 2 shares ~75% nucleotide identity, and intron 3 shares ~60% nucleotide identity (Fig. 2A). *In silico* analysis of intron features revealed a canonical NG|GTNNGT motif at the donor sites for both *RPL22*α and *RPL22***a**, and both canonical and non-canonical acceptor motifs CAG|G/C for *RPL22*α and YAG|Y/G for *RPL22***a**. Four branch sites were predicted for each *RPL22* gene, with the canonical motif CTRAY being more represented (Fig. 2B). Note that the vertical bar represents the exon–intron junction, and Y and R indicate nucleotides with pyrimidine and purine bases, respectively. The length of the predicted polypyrimidine tracts ranged from 9 to 33 nt, and they differed only in intron 1 (Fig. 2A).

**Figure 2.**
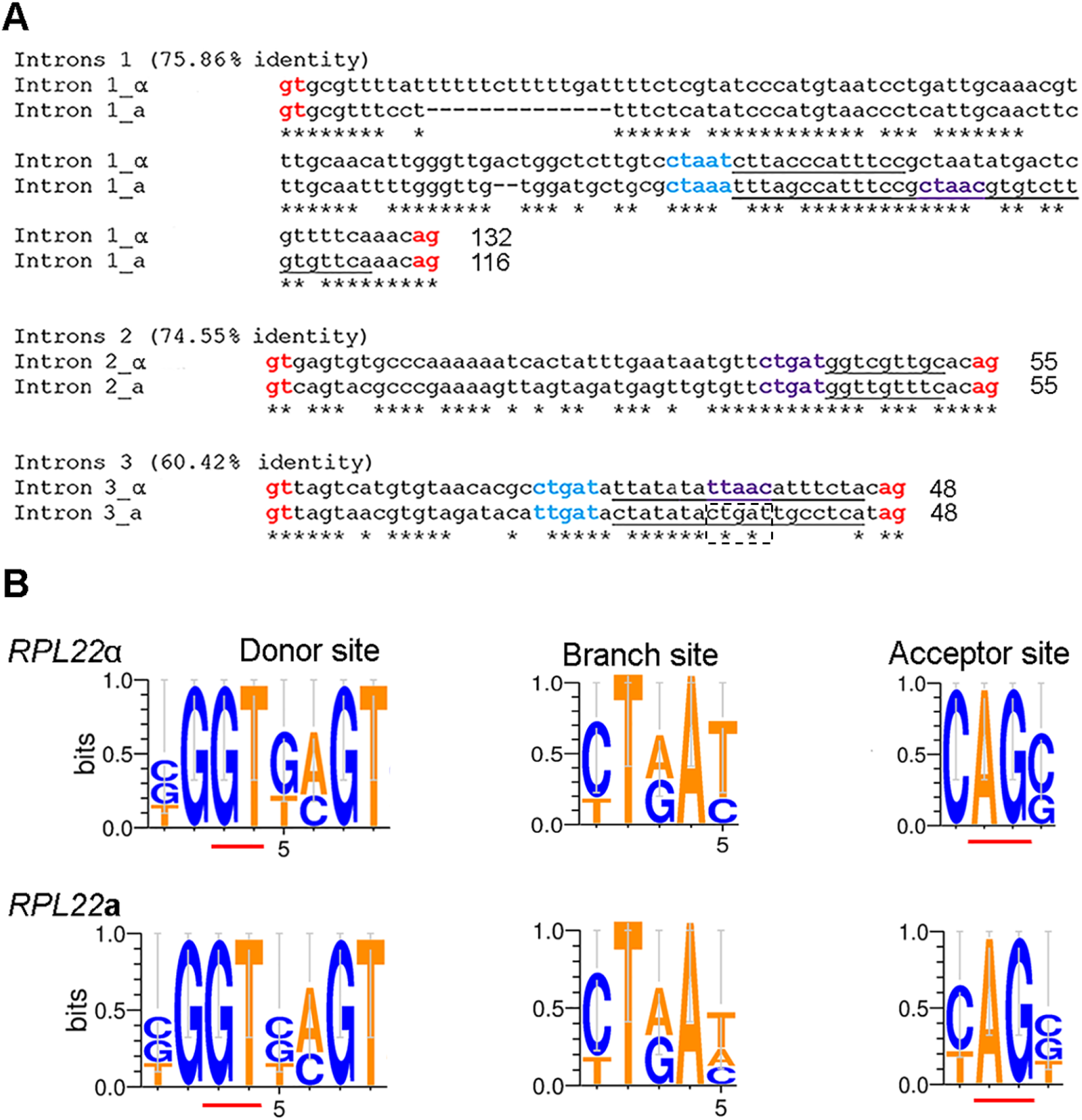
Intron analysis of *C. neoformans RPL22* genes. (A) Alignment of introns 1, 2, and 3 of *RPL22*α and *RPL22***a**. Donor and acceptor splice sites are indicated in red. Branch sites were predicted using a combination of the online software SROOGLE (http://sroogle.tau.ac.il/) (SCHWARTZ *et al.*2009) and SVM-BP finder (http://regulatorygenomics.upf.edu/Software/SVM_BP/): branch sites based on the algorithm of Kol are represented in blue (KOL *et al.* 2005), those based on the algorithm of Schwarts are in purple (SCHWARTZ *et al.* 2008). Manual adjustments were performed based on the websites instructions and score prediction; both algorithms failed to identify a CTRAY canonical branch site of intron 3 of *RPL22***a** (boxed). The polypyrimidine tracts are underlined. The percentage of identity is indicated in parentheses, while the length (in bp) of the introns is indicated at the end on each alignment. (B) Consensus sequences at the 5′ splice site, the branch site, and the 3′ splice site constructed using WebLogo 3.3. bits, binary digits (http://weblogo.threeplusone.com/). Underlined are the canonical donor GT and acceptor AG.

### The *rpl22*a mutant was not complemented by the *RPL22*α allele

We sought to determine whether the Rpl22 *MAT* proteins play a specialized role in *C. neoformans.* The first approach was based on the heterologous expression of *RPL22***a** or *RPL22*α in the heterozygous mutants *RPL22***a**/*rpl22*αΔ (strain GI81) and *RPL22*α*/rpl22***a**Δ (strain GI54). Briefly, the *RPL22* genes including promoters and terminators were cloned into plasmid pSDMA57 for safe haven complementation with the *NEO* selectable marker (ARRAS *et al.* 2015); the empty plasmid served as the control. NAT and NEO double drug resistance coupled with PCR analyses to confirm safe haven integration and lack of plasmid catenation were the basis to select transformants for analysis; in some cases, integration at the safe haven was not achieved, and a heterozygous mutant with ectopic integration of the plasmid was chosen for analysis.

Complementing heterozygous strains were sporulated on MS media, and segregation of the five markers available (*NEO, NAT, MAT, URA5, ADE2*) was determined in the germinated progeny. The criteria for the successful complementation of the mutant phenotype were: 1) presence of both NAT^R^ and NEO^R^ markers in which the progeny’s *MAT* region segregated with the mutated *rpl22*Δ*NAT* allele – this indicates that the ectopic integration of the *RPL22*-*NEO* allele was able to confer viability in progeny that inherited the essential gene *rpl22* deletion 2) absence of only NAT^R^ progeny, because those that inherit the *rpl22*Δ*NAT* allele are inviable; 3) presence of NEO^R^ progeny that, together with progeny sensitive to *NAT* and *NEO*, inherited the *MAT* allele that contains the non-mutated copy of *RPL22*. Progeny that were both *MAT***a** and *MAT*α were expected to be NAT^R^ and either aneuploid or diploid.

Results from the complementation experiments in the heterozygous mutants are summarized in Table 2. For the heterozygous *RPL22*α/*rpl22***a**Δ (GI56), ectopic introduction of a WT copy of *RPL22***a** (strain GI151) was able to restore viability in the progeny, with the generation of three NAT^R^ and NEO^R^ progeny that were *MAT***a** (Fig. 3). Several attempts with both biolistic and electroporation of plasmid pSDMA57 + *RPL22*α in strain GI56 yielded a low number of transformants, a high percentage of plasmid catenation, and lack of integration at the safe haven. Nevertheless, in two independent experiments 2 heterozygous *RPL22*α/*rpl22***a**Δ + *RPL22*α mutants (strains GI102 and GI154) with a single ectopic copy of the plasmid pSDMA57 + *RPL22*α were isolated. Genetic analysis of basidiospores dissected from strain GI102 revealed surprising findings, with no NEO^R^ progeny and two NAT^R^ progeny that had an extra copy of *MAT*α, which includes the *RPL22*α gene and explains their NAT resistance (Fig. 3). Regarding transformant GI154, out of 20 spores that germinated, only one was solely NAT^R^ and three were both NAT^R^ and NEO^R^, and all were both *MAT***a** and *MAT*α. These results indicated that the *RPL22*α gene did not complement the *rpl22***a** mutation. Finally, sporulation of strain GI104 bearing the empty plasmid pSDMA57 also produced one spore that was both *MAT***a** and *MAT*α (Fig. 3).

**Table 2.**
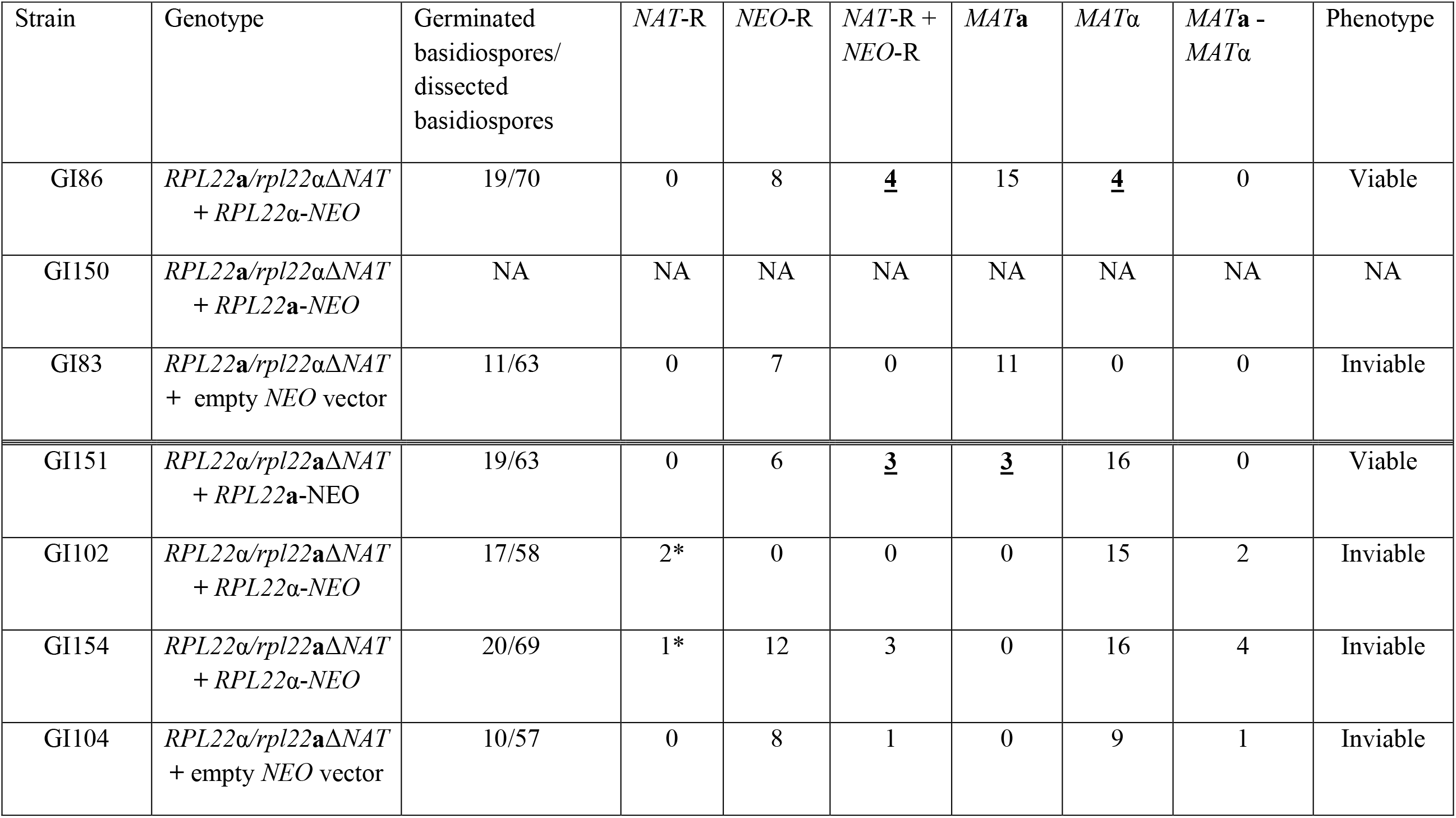
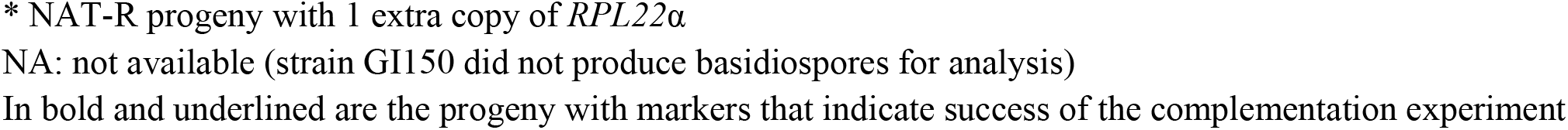
Genetic analysis of *C. neoformans* complementing strains of heterozygous mutants *RPL22***a***/rpl22*αΔ and *RPL22*α*/rpl22***a**Δ

**Figure 3.**
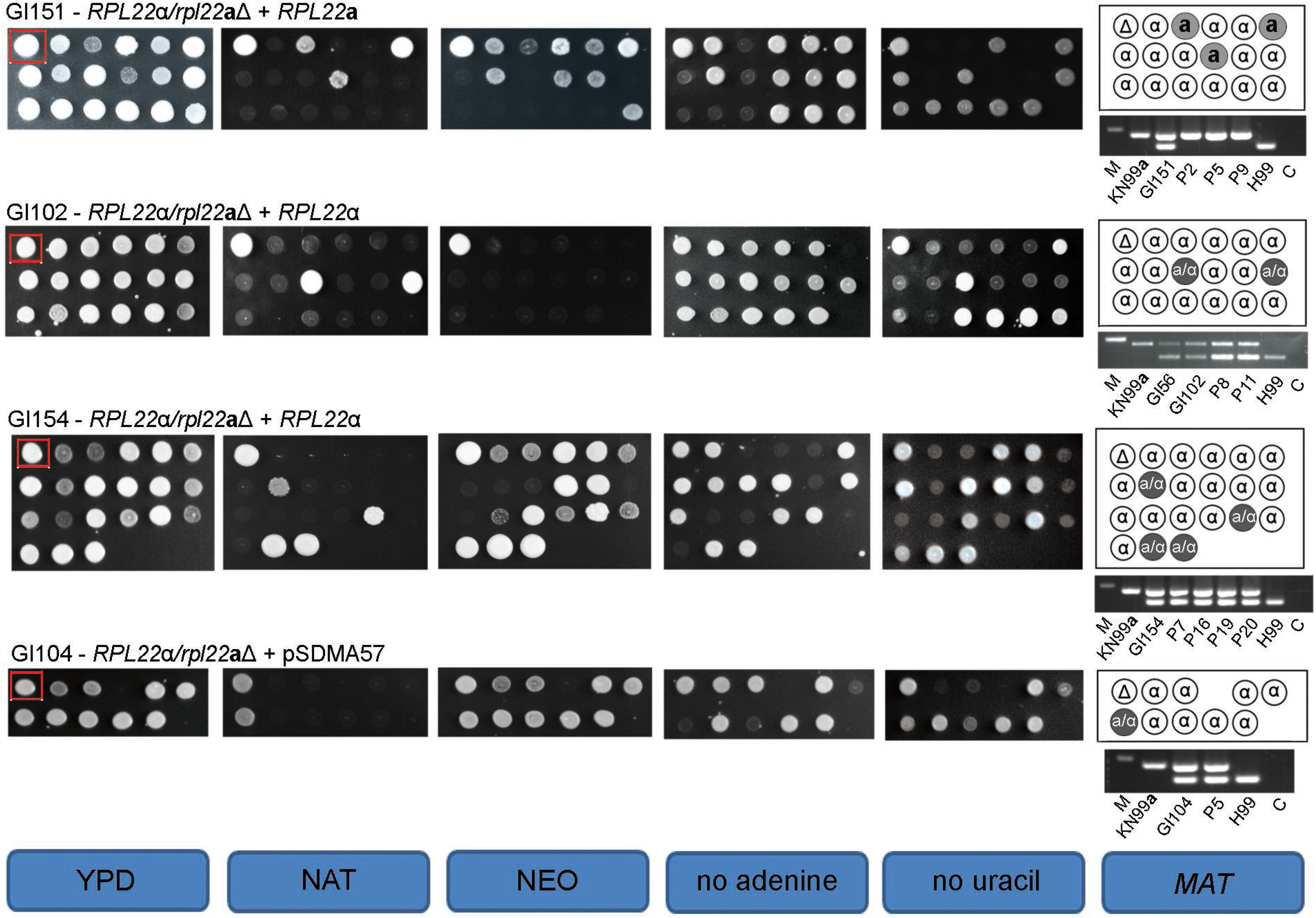
*RPL22***a** complements to restore viability of *rpl22***a** mutants, *RPL22*α does not. Genetic analysis of *C. neoformans* complemented NEO^R^ strains GI151, GI102, GI154 and GI104 derived from heterozygous mutant GI56 (*RPL22*α/*rpl22***a**Δ). The first colony in the top left corner represents the original heterozygous mutant from which the progeny analyzed originate (red boxes). The mating type in NAT^R^ colonies was scored by genetic crosses and by PCR with primers JOHE39201-JOHE39204. For PCR, the WT (H99 *MAT*α and KN99**a***MAT***a**) and the heterozygous strain that was analyzed were used as controls; the NAT^R^ progeny is indicated with “P”, and the number that corresponds to the position in the plate. C = negative control.

Integration of *RPL22*α at the safe haven (strain GI86) restored viability in progeny derived from the heterozygous *RPL22***a**/*rpl22*αΔ mutant, with the generation of four NAT^R^ and NEO^R^ strains that were *MAT*α. For *RPL22***a**/*rpl22*αΔ + *RPL22***a**, none of the transformants obtained had the plasmid integrated at the safe haven. Several transformants having a single ectopic copy of the plasmid were sporulated on MS media, but none was able to form basidiospores for genetic analysis; a representative strain (GI150) is included in Table 2. These results indicate that *RPL22***a**/*rpl22*αΔ + *RPL22***a** complementing strains failed to sporulate and hence we could not assess proper functional complementation through genetic analysis of the meiotic progeny. Finally, as expected introduction of the empty plasmid in *RPL22***a**/*rpl22*αΔ (strain GI83) did not restore progeny viability.

### *RPL22*a expression is regulated by the RNAi pathway

Next, we sought to determine the expression levels of the *RPL22* genes during mating (H99 x KN99**a**) on MS medium in comparison to vegetative growth on YPD agar. While after 24 h of incubation both *RPL22* genes were expressed at a similar level, at 48 h *RPL22*α expression drastically decreased and remained lower than that of *RPL22***a** up to 96 h (Fig. 4A). Expression of *RPL22***a** was also higher than *RPL22*α during vegetative growth on YPD, reflecting the results obtained during mating (Fig. S2). These results indicate that the two *RPL22* alleles are differentially expressed, with *RPL22***a** expression being higher than *RPL22*α during both vegetative growth and mating.

**Figure 4.**
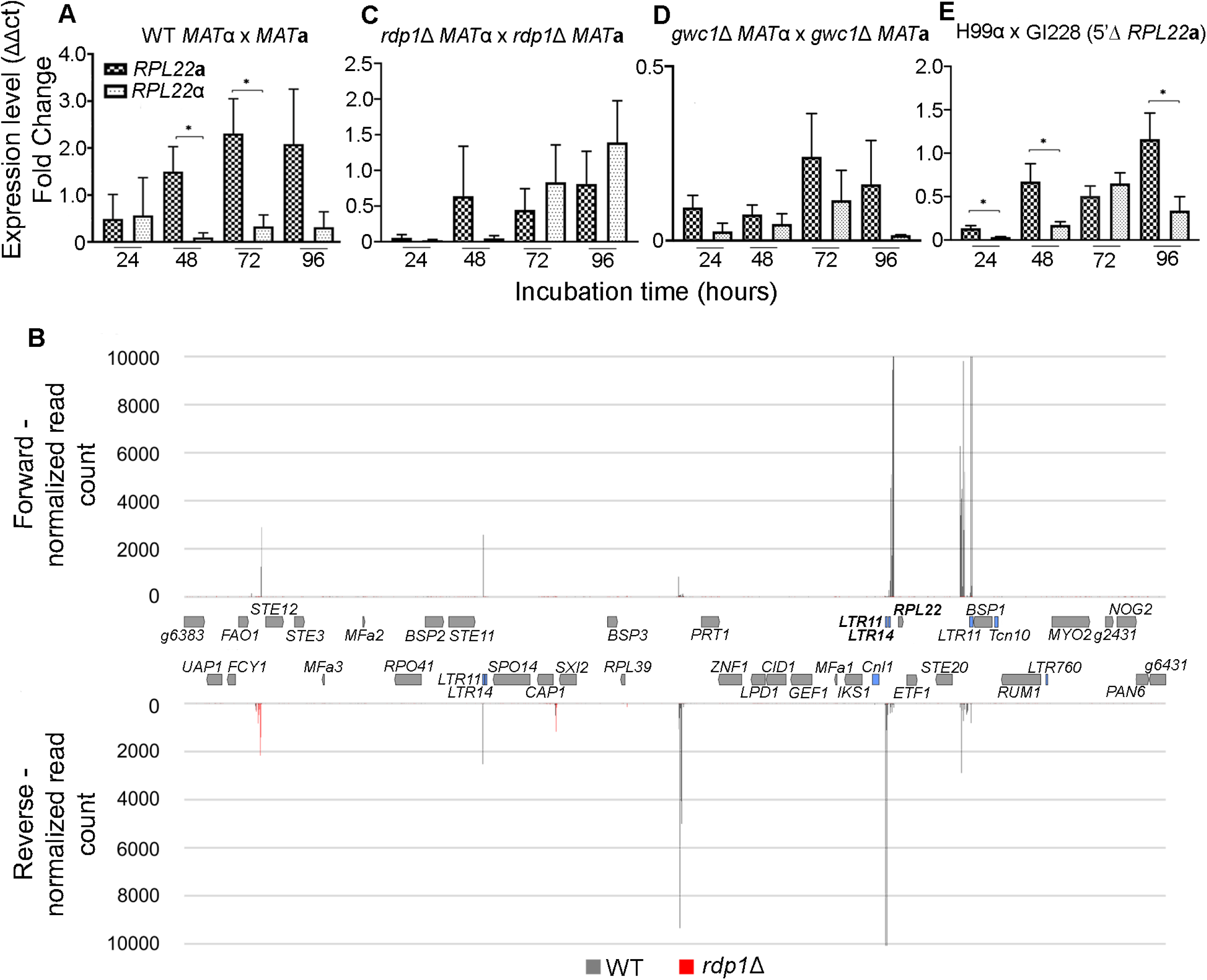
RNAi contributes to control expression of *RPL22***a**. *RPL22*α and *RPL22***a** expression during *C. neoformans* mating between WT H99α x KN99**a** cross (A), *rdp1*Δ and *gwc1*Δ bilateral crosses (C, D), and GI228 x H99 cross (E) for 24, 48, 72 and 96 h of incubation. Ct values were converted to expression level (fold change) through comparison with the endogenous reference *GDP1* (ΔΔct analysis); asterisk indicates p <0.05 for each *RPL22*α and *RPL22***a** comparison for the same day of incubation. Note the different scales on the y axes of the graphs in A - C - D – E. (B) sRNA obtained during H99α x KN99**a** cross (black) and *rdp1*Δ bilateral cross (red) were mapped to the reference *MAT***a** locus of *C. neoformans* (accession number AF542528.2); genes are represented in grey in the middle panel; in blue the *LTR* and transposable elements. *LTR11, LTR14* and *RPL22***a** of interest in this study are indicated in bold.

What are the mechanisms that control *RPL22* expression? Our hypothesis was that inefficiently spliced introns of *RPL22*α could trigger RNA interference (RNAi) through the SCANR complex with subsequent silencing of the gene (DUMESIC *et al.* 2013). Analysis of intron retention (IR) and the splicing pattern of *RPL22*α in several conditions revealed that intron 1 of *RPL22*α is subject to IR, whereas there is minimal IR for introns 2 and 3 (Fig. S3). Mapping small RNA data from an H99α x KN99**a** cross against the *MAT***a** and *MAT*α regions, we found that no reads mapped against the *RPL22* genes, including intronic regions and intron-exon junctions. This observation argues against the hypothesis that there is a direct role of RNAi in governing *RPL22* gene expression.

Interestingly, analysis of the region surrounding the *RPL22* genes revealed that more than 54,989 sRNA reads map to the 2.2 kb region upstream of the *RPL22***a** gene, which includes the *LTR11* and *LTR14* elements, and a candidate long non-coding RNA (lncRNA) predicted based on BLAST analyses. Several sRNA mapping parameters were tested to evaluate whether the sRNA reads map to multiple locations in the genome of *C. neoformans*, allowing us to determine the origin of sRNA reads that were mapped to the 5’ upstream region of *RPL22***a**. We found that the majority of the reads originated from the 5′ upstream region of the *RPL22***a** gene, while others originated from regions of the genome distant from *MAT* corresponding to the lncRNAs CNAG_12037 and CNAG_13142, and the region upstream of the lncRNA CNAG_12435.

Analysis of the region upstream of the *RPL22***a** gene in an *rdp1*Δ *MAT*α x *rdp1*Δ *MAT***a** bilateral cross revealed a drastic reduction in sRNA reads (Fig. 4B), consistent with a role for RNAi in governing these sRNA. We then performed RT-qPCR of the *rdp1*Δ x *rdp1*Δ bilateral cross, and found that *RPL22***a** expression was lower than *RPL22*α expression at 72 h and 96 h of incubation (Fig. 4C). Compared to a WT H99α x KN99**a** cross, *RPL22***a** expression was overall lower whereas that of *RPL22*α was higher (Fig. S4A). Increased expression of *RPL22*α in the *rdp1*Δ mutant bilateral cross corroborates previous findings (WANG *et al.* 2010), although *RPL22*α expression seems to be indirectly regulated by RNAi because no sRNA map to the regions surrounding the *RPL22*α gene (Fig S5).

We also performed RT-qPCR during a bilateral cross between mutants for the SCANR complex component Gwc1, and found strong downregulation [fold change (FC) < 0.5)] of both *RPL22***a** and *RPL22*α (Fig. 4D; Fig. S4B). Lastly, we used the recently-developed CRISPR/Cas9 technology to accurately delete the *MAT* region in *C. neoformans* KN99**a** that is upstream of *RPL22***a** and that is targeted by the abundant sRNA. The strain generated was named GI288 (5ʹΔ *RPL22***a**), and a schematic representation of the strategy is shown in Fig. S6A. RT-qPCR expression analysis during mating revealed that *RPL22***a** expression remained higher than *RPL22*α, except at 72 h of incubation (Fig. 4E). As compared to *RPL22* expression during the WT cross, *RPL22***a** was strongly downregulated in the GI228 (5ʹΔ *RPL22***a**) x H99α cross at 48 h and 72 h of incubation, while expression of *RPL22*α remained unchanged (Fig. S6B). Interestingly, expression of *RPL22***a** during the GI288 (5ʹΔ *RPL22***a**) x H99α cross mirrors that of *RPL22***a** during the *rdp1*Δ bilateral cross, in accord with an RNAi-dependent mechanism regulating *RPL22***a** expression (Fig. S6C). Despite the downregulation of *RPL22***a**, the GI228 mutant strain does not display any morphological defect during vegetative growth, and its ability to mate with H99 and generate viable progeny was not compromised; as expected, genetic analysis of progeny derived from the GI228 x H99 cross showed co-segregation of *NAT* with *MAT***a** (data not shown).

### *RPL22* exchange allele strains exhibit sexual reproduction defects

We next sought to determine whether functional complementation of the *RPL22* genes could be achieved by replacing either of them with the opposite *RPL22* allele at the native locus within the *MAT* loci. To this end, we generated exchange alleles of *RPL22*α and *RPL22***a** by means of CRISPR/Cas9, whose use was critical due to suppressed recombination within *MAT*. Because the *RPL22* genes share a high level of identity (83%) (Fig. S1), specificity of the gene replacement was achieved by designing two specific guide RNA (gRNA) molecules that determined the sites for homologous recombination. Transformation was performed by electroporation with the simultaneous introduction of the three gRNAs [at the 5ʹ and 3ʹ of the *RPL22* gene, and one for the Safe Haven 2 (SH2)], the homology-directed repair (HDR) *RPL22* gene, and selectable markers *NAT* or *NEO,* which were introduced in the Safe Haven 2 (SH2) region to avoid unnecessary ectopic mutations that could interfere with the resulting phenotype. Recipients for transformations were the most isogenic strains available for *C. neoformans* serotype A, H99 (*MAT*α) and KN99**a** (*MAT***a**) (NIELSEN *et al.* 2003; JANBON *et al.* 2014; FRIEDMAN *et al.* 2018).

In two independent transformation attempts, precise gene replacement of *RPL22*α with *RPL22***a** at its native location within the *MAT*α locus of H99 and correct integration of *NAT* in the SH2 were readily obtained. This resulted in the generation of *rpl22*α∷*RPL22***a***SH2*∷*NAT* mutant strains (YFF92) that differ from their parental strain at only the *RPL22* gene. A schematic representation of the exchange strains generated is shown in Figure S7. Conversely, via the same strategy (i.e. two gRNA at the 5ʹ and 3ʹ of the *RPL22*) an *RPL22***a** exchange allele strain could not be isolated. Because the *RPL22* genes differ in only 5 amino acids that are located in the N-terminal region (Fig. S1, S7), to replace *RPL22***a** with the Rpl22α coding gene a different strategy based on CRISPR/Cas9 was employed. In this approach we generated a chimeric *RPL22*α (c*RPL22*α) HDR template, which consisted of the N-terminus of *RPL22*α fused with the C-terminus of *RPL22***a**, and this was introduced into *C. neoformans* strain KN99**a** together with new gRNAs designed to target the 5ʹ region of *RPL22***a** (Fig 5A; Fig. S7). Several independent *rpl22***a**∷*RPL22*α^N^-*RPL22***a**^C^*SH2*∷*NEO* exchange strains were obtained, and one (strain YFF113) was chosen for further experiments. Allele exchange strains YFF92 and YFF113 were used in unilateral and bilateral crosses to evaluate both their *MAT*-specificity and the phenotypic consequence due to the absence of one *RPL22* gene. In the presence of only *RPL22***a** (cross KN99**a** x YFF92α) or *RPL22*α (cross H99α x YFF113**a**), the strains displayed no altered morphology and had WT mating ability, spore germination, independent segregation of the markers, and uniparental inheritance of mitochondria (Fig. 5B; File S2; Table 3). These results indicate that the absence of one Rpl22 allele does not affect mating efficiency or meiosis when the other allele is present in the native location within *MAT*. Interestingly, RT-qPCR revealed a higher level of expression of the chimeric c*RPL22*α gene in the KN99**a** background compared to that of *RPL22***a** in the H99 background at 72 h and 96 h of incubation, hence displaying an opposite trend compared to the WT cross (Fig. 5D; Fig. S8A).

**Table 3.** Genetic analysis of crosses between allele exchange strains

| Cross | Spores dissected | Spores germinated | *MAT*α-*RPL22*α |  | *MAT*a - *RPL22*a |  | *MAT*a - *MAT*α | Mitochondria from *MAT*a | Mitochondria from *MAT*α |  |
| --- | --- | --- | --- | --- | --- | --- | --- | --- | --- | --- |
| H99 x KN99a | 60 | 27 | 11 |  | 14 |  | 2 | 27 | 0 |  |
| YFF92 x KN99a | Spores dissected | Spores germinated | *MAT*α-*RPL22*a | *MAT*a - *RPL22*a | *MAT*a - *MAT*α | NAT-R progeny (*MAT*a; *MAT*α; *MAT*a/α) |  | Mitochondria from *MAT*a | Mitochondria from *MAT*α |  |
|  | 60 | 18 | 9 | 6 | 3 | 7 (4; 2; 1) |  | 18 | 0 |  |
| YFF113 x H99 | Spores dissected | Spores germinated | *MAT*a - *cRPL22*α | *MAT*α-*RPL22*α | *MAT*a - *MAT*α | NEO-R progeny (*MAT*a; *MAT*α; *MAT*a/α) |  | Mitochondria from *MAT*a | Mitochondria from *MAT*α |  |
|  | 62 | 49 | 27 | 19 | 3 | 31 (16; 12; 3) |  | 49 | 0 |  |
| YFF116 x H99 | Spores dissected | Spores germinated | *MAT*a-*RPL22*α | *MAT*α-*RPL22*α | *MAT*a - *MAT*α | NEO-R progeny (*MAT*a; *MAT*α; *MAT*a/α) |  | Mitochondria from *MAT*a | Mitochondria from *MAT*α |  |
|  | 66 | 43 | 23 | 17 | 3 | 23 (10; 10; 3) |  | 43 | 0 |  |
| YFF92 x YFF113 | Spores dissec. | Spores germ. | *MAT*a-*cRPL22*α | *MAT*α-*RPL22*a | *MAT*a - *MAT*α | NEO-R (*MAT*a; *MAT*α; *MAT*a/α) | NAT-R (*MAT*a; *MAT*α; *MAT*a/α) | NEO-R NAT-R | Mito. *MAT*a | Mito. *MAT*a |
|  | 63 | 28 | 14 | 13 | 1 | 14 (9; 5; 0) | 14 (6; 7; 0) | 0 | 28 | 0 |
| YFF92 x YFF116 | Spores dissec. | Spores germ. | *MAT*a-*RPL22*α | *MAT*α-*RPL22*a | *MAT*a - *MAT*α | NEO-R (*MAT*a; *MAT*α; *MAT*a/α) | NAT-R (*MAT*a; *MAT*α; *MAT*a/α) | NEO-R NAT-R | Mitoc. *MAT*a | Mitoc. *MAT*a |
|  | 58 | 7 | 3 | 0 | 4 | 1 (1; 0; 0) | 4 (0; 0; 4) | 0 | NA | NA |

NA: not available

**Figure 5.**
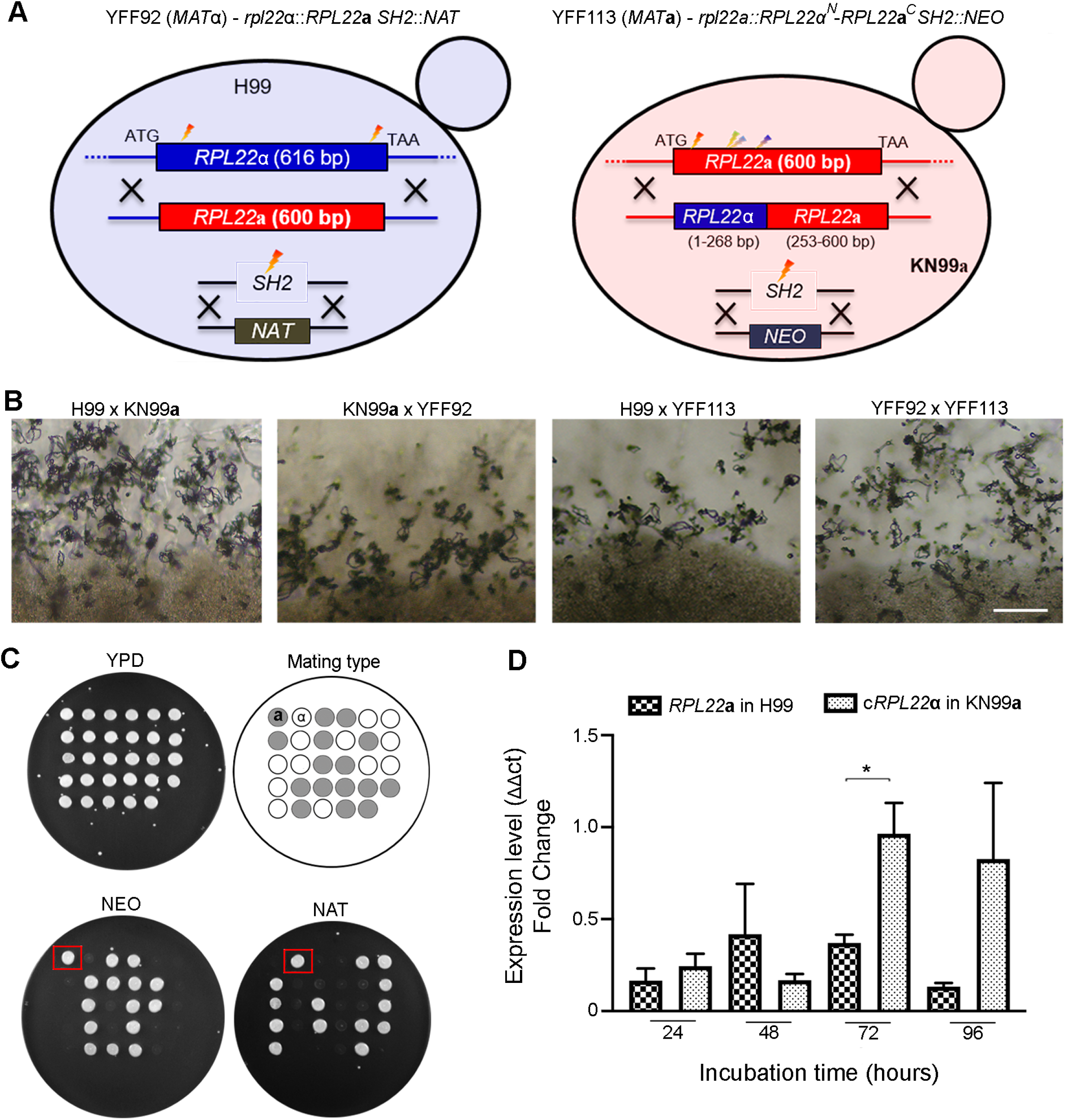
Construction, analysis and sexual reproduction of *RPL22* exchange strains. (A) Schematic representation of the generation of the *C. neoformans RPL22* exchange strains YFF92 and YF113. Lightning bolts in different colors denote different gRNA targeting sites. (B) Mating phenotypes during crosses between H99α x KN99**a**, and YFF92 x KN99**a**, H99α x YFF113, and YFF92 x YFF113. The scale bar is 100 μm. (C) Genetic analysis of progeny obtained from the YFF92 x YFF113 bilateral cross; note the expected 1:1 segregation of the *SH2*∷*NAT* and *SH2*∷*NEO* markers in progeny produced from YFF92 x YFF113 cross, and the independent segregation of the *MAT* loci, with *MAT***a** indicated in grey and *MAT*α indicated in white. (D) RT-qPCR of *RPL22***a** and c*RPL22*α expression during the YFF92 x YFF113 cross. Asterisk indicates p<0.05 for each c*RPL22*α and *RPL22***a** comparison for the same day of incubation.

Because the approach utilized to generate strain YFF113 was successful, we employed the same gRNA to replace the *RPL22***a** gene with a native copy of the *RPL22*α gene. After several unsuccessful attempts, we obtained one *rpl22***a**∷*RPL22*α strain (YFF116) with the *NEO* marker integrated ectopically in the genome, and not in the *SH2* locus as planned (Fig. 6A; Fig. S7). Similar to findings presented above, in the presence of only the *RPL22*α gene (cross H99α x YFF116**a**) no morphological and genetic defects were observed (Fig. 6C; File S2; Table 3). Remarkably, bilateral crosses of strains with exchanged *RPL22* genes (YFF92α x YFF116**a**) exhibited a high percentage of irregular basidia (Fig. 6D). High resolution scanning electron microscopy revealed that the majority of the basidia had no basidiospores, while others had a morphology defect (Figure 6E-F; File S3), and a low number had irregular basidiospore chains collapsed on the basidia (Figure 6G; File S3). The formation of clamp connections was not affected (Figure 6H; File S3). Basidiospores germinated from cross YFF92α x YFF116**a** displayed a low germination rate (12%) and irregular segregation of the meiotic markers, with 3 progeny being *MAT***a***-RPL22*α, no *RPL22***a***-MAT*α progeny, and 4 progeny being both *MAT*α and *MAT***a**, hence aneuploid or diploid (Table 3). RT-qPCR analysis during YFF92 x YFF116 mating revealed low expression (FC < 1) of both *RPL22* genes from 24 h to 96 h (Fig. 6B; Fig. S8B).

**Figure 6.**
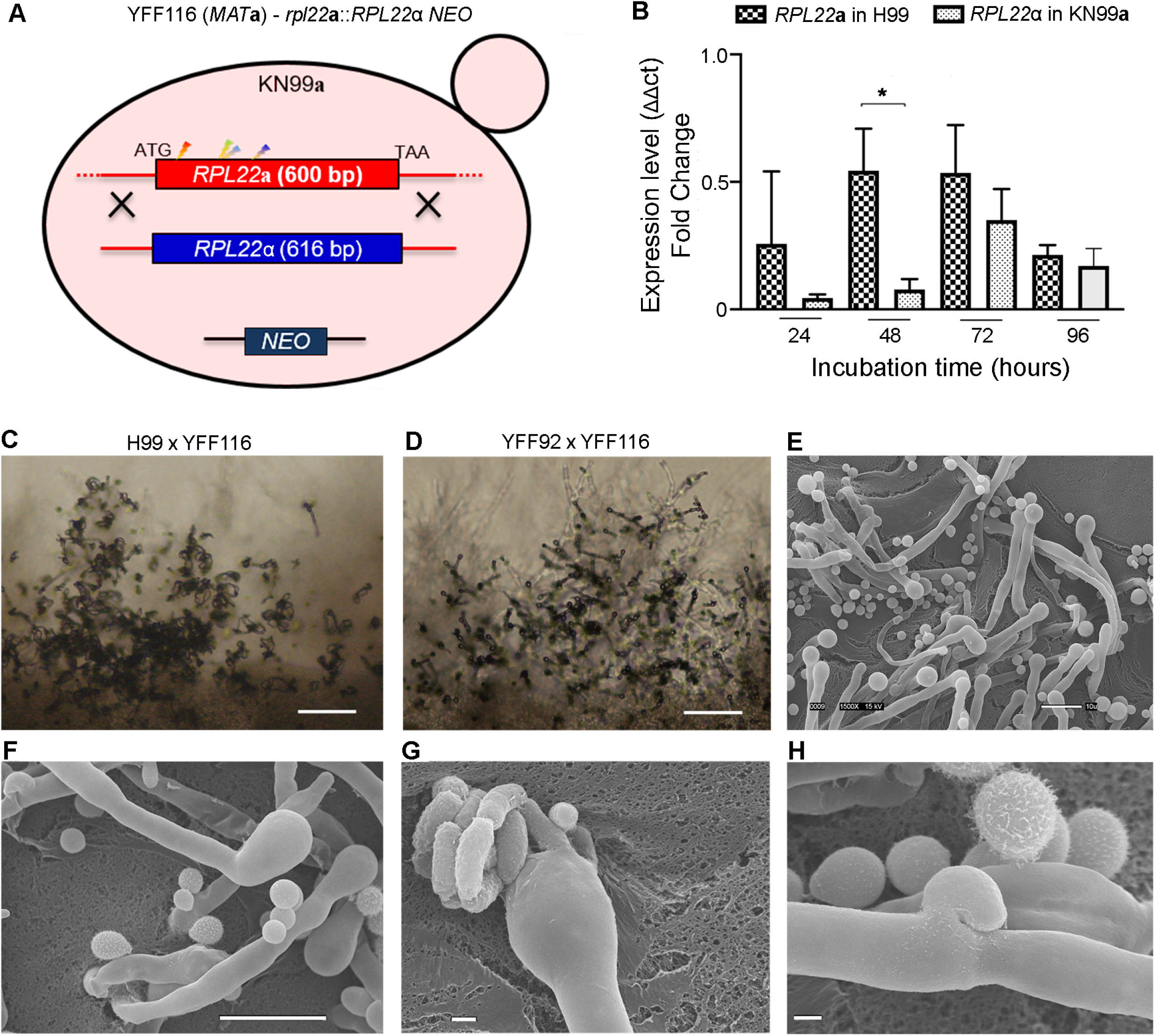
Reciprocal *RPL22* exchange strains exhibit defects in sexual reproduction. (A) Schematic representation of the generation of the *C. neoformans RPL22* swapped strain YFF116. Lightning bolts in different colors denote different gRNA targeting sites. (B) RT-qPCR of *RPL22***a** and *RPL22*α expression during YFF92 x YFF116 cross; asterisk indicates p<0.05 for each c*RPL22*α and *RPL22***a** comparison for the same day of incubation. (C - D) Mating phenotypes during crosses between the cross H99 x YFF116, and YFF92 x YFF116. The scale bar is 100 μm. (E) Scanning electron microscopy of sexual structures of the YFF92 x YFF116 cross; note that the majority of the basidia are bald with no basidiospore chains. The scale bar is 10 μm. (F) Scanning electron microscopy of an irregular basidium produced by the YFF92 x YFF116 cross. The scale bar is 10 μm. (G) Scanning electron microscopy of a basidium with a collapsed chain of basidiospores produced by the YFF92 x YFF116 cross. The scale bar is 1 μm. (H) Scanning electron microscopy of a regular unfused clump connection produced by the YFF92 x YFF116 cross. The scale bar is 1 μm.

To confirm that this defect was due to the *rpl22***a**∷*RPL22*α exchange allele and to exclude any influence of the *NEO* marker, 3 NEO^S^ *MAT***a** F1 progeny (SEC876, progeny 7; SEC884, progeny 15; SEC889, progeny 20) obtained from the H99α x YFF116**a** cross were backcrossed with both H99α and YFF92α. Corroborating the results obtained with strain YFF116, progeny SEC876, SEC884, and SEC889 displayed normal mating with H99α, but when crossed with the the exchange allele strain YFF92 all three exhibited morphological defects remarkably similar to the YFF116 parent (Fig. S10B).

### Phenotypic analysis of *rpl22* mutant strains

Heterozygous deletion mutants and exchange strains (Table S1) were tested for altered phenotypic traits (see Materials and Methods for details). Of the 28 stresses tested, few phenotypic differences among the isolates were observed on FLC, YP + NaCl, caffeine, 39°C and 4°C (Fig. S9). Exchange strain YFF116 (*rpl22***a**∷*RPL22*α *NEO*) displayed sensitivity to hydroxyurea compared to its parental strain KN99**a** (Fig. S9), but genetic analysis of the markers revealed that this was due to the ectopic integration of NEO (Fig. S10A).

## Discussion

A previous study reported that the *MAT* locus of *C. neoformans* contains five genes (*RPL39, RPL22, MYO2, RPO41, PRT1*) that encode proteins required for viability (FRASER *et al.* 2004). The essential nature of these genes was inferred based on the inability to mutate them in a haploid strain of *C. neoformans.* In addition to *C. neoformans*, *Candida albicans* is the only other fungus known to encode essential genes within the *MAT* locus (HULL *et al.* 2000; SRIKANTHA *et al.* 2012). It is likely that one function of *MAT*-essential genes is to constrain recombination and serve as a genetic buffer constraining loss of portions of the *MAT* locus, although they might also serve other functions related to development, sexual reproduction, and virulence.

Here the functions of the α and **a***MAT* specific alleles encoding the *C. neoformans* ribosomal proteins Rpl22 and Rpl39 were characterized. Given suppressed recombination within the *C. neoformans MAT* loci, mutation of the *RPL22* and *RPL39* genes was challenging. Heterozygous mutants for these genes were successfully generated through the combined use of a biolistic split marker approach and the compound W7 to inhibit non-homologous end joining and enhance homologous recombination (FU *et al.* 2006; ARRAS AND FRASER 2016). Through deletions in the *C. neoformans* **a**/α diploid strain AI187 and Mendelian analysis of recombinant F1 progeny obtained following sexual reproduction and spore dissection, we demonstrated that both the α and **a** alleles of the *RPL22* and *RPL39* genes are essential (Table 1; Fig. 1). Conversely, in *S. cerevisiae* the *RPL22* and *RPL39* orthologs are not required for viability (STEFFEN *et al.* 2012; KIM AND STRICH 2016), indicating evolutionary divergence of essential ribosomal genes between Ascomycetous and Basidiomycetous yeasts.

The ribosome was thought to be a constant, conserved, uniform protein translation machine. Recent studies have revealed novel and unexpected findings for the ribosome, in particular complex heterogeneity and specialized activity that confers regulatory control in gene expression (WARNER AND MCINTOSH 2009; NARLA AND EBERT 2010; XUE AND BARNA 2012). While in mammals ribosomal proteins are encoded by single genes, in yeasts, plants, and flies, ribosomal proteins are encoded by several genes. A remarkable example is the model yeast *S. cerevisiae*, in which, following a genome duplication event, 59 of the 78 ribosomal proteins are encoded by two retained gene copies and share high sequence similarity, but are in most cases not functionally redundant, and have been found to play specialized functions [(XUE AND BARNA 2012) and references within it]. Specialized ribosomes have been identified also in plants, flies, zebrafish, and mice (KOMILI *et al.* 2007; MCINTOSH AND WARNER 2007; XUE AND BARNA 2012), but it is not known whether they exist in microbial pathogens.

A number of studies have converged to reveal diverse and specialized roles for the Rpl22 ribosomal paralogs in yeasts and vertebrates. In *S. cerevisiae*, haploid *rpl22*a mutants are cold sensitive, and display reduced invasive growth and a longer doubling time compared to *rpl22*b (STEFFEN *et al.* 2012; KIM AND STRICH 2016). Moreover, *rpl22*a mutations perturb bud site selection and cause random budding, while *rpl22*b mutations do not, and overexpression of *RPL22*B in *rpl22*a mutants fails to restore bud site selection (KOMILI *et al.* 2007). Vertebrates also express Rpl22 paralogs, called Rpl22 and Rpl22-like1 (RPL22-l1). Mice lacking *RPL22* are viable and have specific αβ T-cell developmental defects, likely attributable to compensation by Rpl22- l1 in other tissues (ANDERSON *et al.* 2007). Recent studies in both yeast and mammals suggest an extraribosomal role for Rpl22 paralogs in binding target mRNAs and regulating their expression (GABUNILAS AND CHANFREAU 2016; ZHANG *et al.* 2017; ABRHAMOVA *et al.* 2018). From an evolutionary viewpoint, it is important to highlight that in *C. neoformans* the *RPL22* gene is present as a single copy and it is not the result of a genome duplication event in contrast to *S. cerevisiae*. Instead, the *C. neoformans RPL22* gene was relocated to within the *MAT* locus concurrent with the transition from tetrapolar to bipolar, and because of the suppressed recombination in this region, the two *RPL22*α and *RPL22***a** alleles underwent a different evolutionary trajectory that generated differences between them (COELHO *et al.* 2017; SUN *et al.* 2017). Therefore, technically they are alleles rather than paralogs. In this study we sought to determine whether the Rpl22 *MAT* alleles play any specialized role in *C. neoformans.*

We found that *RPL22*α was not able to complement the essential phenotype due to mutations of *RPL22***a**, whereas ectopic introduction of *RPL22***a** in *RPL22***a**/*rpl22*αΔ resulted in sporulation failure. Conversely, viability was restored in progeny derived from heterozygous *RPL22***a**/*rpl22*αΔ + *RPL22*α and *RPL22*α/*rpl22***a**Δ + *RPL22***a** strains (Fig. 3; Table 2). In a parallel approach we ectopically inserted an *RPL22* allele into a *C. neoformans* haploid strain, and then attempted to mutate the native opposite *MAT* copy. Also in this case deletion mutants could not be recovered, further supporting the observation of a failure of complementation between the two *C. neoformans RPL22* alleles (data not shown). We further demonstrate that during both mitotic growth and sexual reproduction *RPL22***a** expression is much higher than *RPL22*α (Fig. 4A, Fig. S2). Considering that the heterozygous mutants *RPL22***a**/*rpl22*αΔ and *RPL22*α/*rpl22***a**Δ are viable, we propose two possible models to explain the lack of complementation. The first is a model involving an expression effect in which differential expression levels of the two *RPL22* alleles at ectopic locations might hamper functional complementation; the second is a model involving position effect in which each *RPL22* allele has to be in its own *MAT* locus.

We determined that an RNAi-mediated mechanism regulates *RPL22***a** expression and that it involves a region located upstream of *RPL22***a** that includes the *LTR11* and *LTR14* elements and a predicted lncRNA; when this upstream region is silenced by sRNA, expression of *RPL22***a** is enhanced. Conversely, in the absence of sRNA (i.e. in an RNAi mutant background), or when the sRNA-targeted region was deleted (Fig. S6A), *RPL22***a** expression was strongly decreased (Fig. 4 B – E, Fig. S4, Fig. S6). This is a novel and intriguing epigenetic mechanism of gene expression regulation within the *MAT***a** locus of *C. neoformans*. Examples of LTR elements silenced by sRNA have been described also in plants and mammals as a mechanism of genome protection (ŠURBANOVSKI *et al.* 2016; MARTINEZ *et al.* 2017; SCHORN *et al.* 2017; MARTINEZ 2018). There are also other locations within the *MAT***a** locus that are characterized by *LTR* elements that are also robustly targeted by sRNA in an RNAi-dependent manner (Fig. 4B), and future studies will elucidate their impact on the gene expression, mating, and genome stability.

While we found differential expression between the *RPL22***a** and *RPL22*α genes, and have identified epigenetic regulation of *RPL22***a** expression, the approaches employed did not enable us to define whether the Rpl22 alleles play specific cellular roles. We then applied a newly developed CRISPR approach to generate haploid isogenic *C. neoformans* strains exchanging the *MAT RPL22* genes: *MAT*α-*RPL22***a** (strain YFF92α) and *MAT***a**-*RPL22*α (strain YFF116**a**) (Fig. S7). Unilateral crosses involving *MAT*α-*RPL22***a** x KN99**a**, and H99α x *MAT***a**-*RPL22*α, exhibited sexual reproduction features similar to the wild type cross H99α x KN99**a**, including dikaryotic hyphae, clamp connections, basidia, and basidiospore chains (Figs. 5-6, File S2). Conversely, the bilateral cross *MAT*α-*RPL22***a** x *MAT***a**-*RPL22*α (cross YFF92α x YFF116**a**) produced regular hyphae and clamp connections, but with irregular basidia and few or no spores that were characterized by a low germination and viability following microdissection, suggesting a defect in nuclear fusion, meiosis, or sporulation (Fig. 6, File S3). This is likely due to the drastic reduction of both *RPL22***a** and *RPL22*α expression (Fig. 6B).

Lastly, we have also generated a chimeric c*RPL22*α-*MAT***a** (YFF113) exchange strain of *C. neoformans* to initially test the phenotypic consequences of exchanging Rpl22, with a focus on the 5 amino acid differences located in the N-terminal region of the protein (Fig. 5; Fig. S7). While this exchange strain does not display any morphological or phenotypic defects (Fig. 5; Fig. S9), its analysis turned out to be of interest with respect to the mechanisms of regulation of *RPL22*α. The strains *MAT***a**-c*RPL22*α (YFF113) and *MAT***a**-*RPL22*α (YFF116) both encode an Rpl22α protein, yet they display very different *RPL22* expression patterns and distinct phenotypes (Fig. S8C, Fig S9). In *S. cerevisiae* introns play a crucial role for *RPL22* expression, with Rpl22 playing an extraribosomal role in inhibiting the splicing of the *RPL22B* pre-mRNA transcript through direct binding of its intron (GABUNILAS AND CHANFREAU 2016; ABRHAMOVA *et al.* 2018). Moreover, this mechanism of autoregulation seems to be conserved also in *Kluyveromyces lactis*, a Saccharomycotina species that did not undergo the whole genome duplication event and retains only one copy of the *RPL22* gene (SCANNELL et al. 2007). Similar mechanisms might operate to control expression of *C. neoformans RPL22*α. Considering the high expression of the chimeric c*RPL22*α but not *RPL22*α (Fig. S8C), one could hypothesize that their different introns could potentially have a regulatory role in *RPL22*α expression. The two Rpl22α-coding genes in strains *MAT***a**-c*RPL22*α (YFF113) and *MAT***a**-*RPL22*α (YFF116) differ only in the 3ʹ region, which for strain YFF113 is from *RPL22***a** and includes introns 2 and 3. Intron 1, which is the largest and most divergent between the *RPL22* genes (Fig. 2; Figs. S7, S11), is the same in the *RPL22*α and c*RPL22*α allele and can therefore be excluded. Introns 2 of *RPL22***a** and *RPL22*α share high sequence similarities and have the same intronic features (Fig. 2; Fig. S11). Introns 3 share lower sequence similarities, and the main differences are found in pre-mRNA secondary structure and nucleotide composition in the region between the branch site location and the 3′ acceptor site (Fig. 2; Fig S11), which is known to affect splicing and gene expression (GAHURA *et al.* 2011; PLASS *et al.* 2012; ZAFRIR AND TULLER 2015). Furthermore, another issue could also be that the canonical branch site of intron 3 of *RPL22*α might be too close to the donor site with inhibition of the lariat formation, while intron 3 of *RPL22***a** has a possible more distal canonical branch site (Fig. 2). Based on these observations, we speculate that intron 3 of *RPL22*α might be a candidate for regulatory function.

Our findings, such as the absence of morphological defects of heterozygous *RPL22/rpl22* mutants, the lack of cross complementation between the *RPL22* alleles, and the morphological and genetic defect of exchange strain MAT**a-***RPL22*α, may support a model in which the two *RPL22 MAT* essential genes operate as a type of imprinting system to ensure fidelity of sexual reproduction to enforce coordinate segregation of the opposite *MAT* nuclei in the dikaryotic hyphae.

## ACKNOWLEDGMENTS

We thank Alexander Idnurm for critical comments on the manuscript.

